# Simulations of Lévy walk

**DOI:** 10.1101/2020.10.09.332775

**Authors:** Venkat Abhignan, Sinduja Rajadurai

## Abstract

We simulate stable distributions to study the ideal movement pattern for the spread of a virus using autonomous carrier. We observe Lévy walks to be the most ideal way to spread and further study how the parameters in Lévy distribution affects the spread.

## 1 Introduction

Biological systems cannot be described only in the aspects of probabilistic distributions. However in the interest of studying interactions and common behaviours we consider systems to imitate similar laws at different scales [9]. Stable distributions have scale invariant behaviour, where linear combination of independent and identical distributions (IID’s) with finite mean and variance leads to a normal distribution (ND) [8]. Similarly when we consider rescaled and reordered sum of IID’s with non-finite variance it may converge to a Lévy distribution (LD). Though ND’s are pervasive in most systems, LD’s are found in biological systems mostly associated with optimal foraging behaviour [9, 14] and human mobility. Most literature supports the statistical similarity between human mobility through different modes of transportation and Lévy distribution [1, 6, 16] but in some instances a log-normal distribution was also observed [20]. LD comprises of Lévy walk (LW) where multiple short steps are taken with long steps in-between, whereas the ND comprises of Brownian walk (BW) where multiple similar steps are taken. Observing the spread of present virus [3] we considered studying simulations of a simple model using stable distributions. It can be observed that spread takes a Lévy like walk which seems to happen in different distance scales. Initially spread across different continents taking long steps followed by multiple short steps within the continent, then again long steps across different countries within the continent and followed by multiple short steps within the country, then different states within the country and so on, the distance scales keep changing but the behaviour nearly is the same. This similar behaviour can perhaps be observed in the recently available data of COVID-19 taken up to 10th November 2020 [3]. Fig 1(a) shows the data of daily new cases in different countries and Fig 2(a) shows the data of daily new cases per 100,000 people in different states within United States. We can observe that different distributions of these countries and states are characterized by a initial peak followed by a decline or flattening of the curve and then followed by a higher peak than the initial. We assume that the macroscopic behaviour of such a system is nearly the same which can be a continent, or a country within the continent, or a state within the country. Independent of the microscopic interactions which influence the local behaviour such as environmental factors, demographics, government policies implemented to control the spread and various other factors that are different in each scale and each system. We qualitatively concern our study only on which movement pattern is ideal for the spread of a virus on a macroscopic scale, without the consideration of such influence on the microscopic scales. We implement simple parameters which depend only on the stable distributions to observe simulations. These simulations are based on the logic of steering behaviour developed by Craig Reynolds [15] and Daniel Shiffman [17].

**Figure 1:**
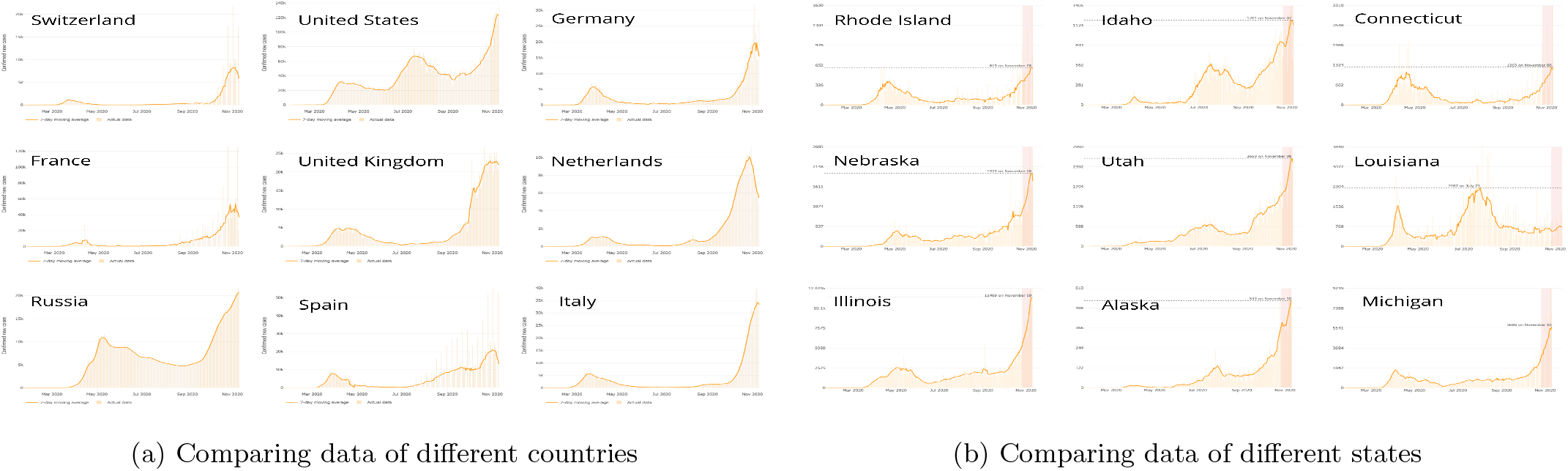
COVID-19 data

**Figure 2:**
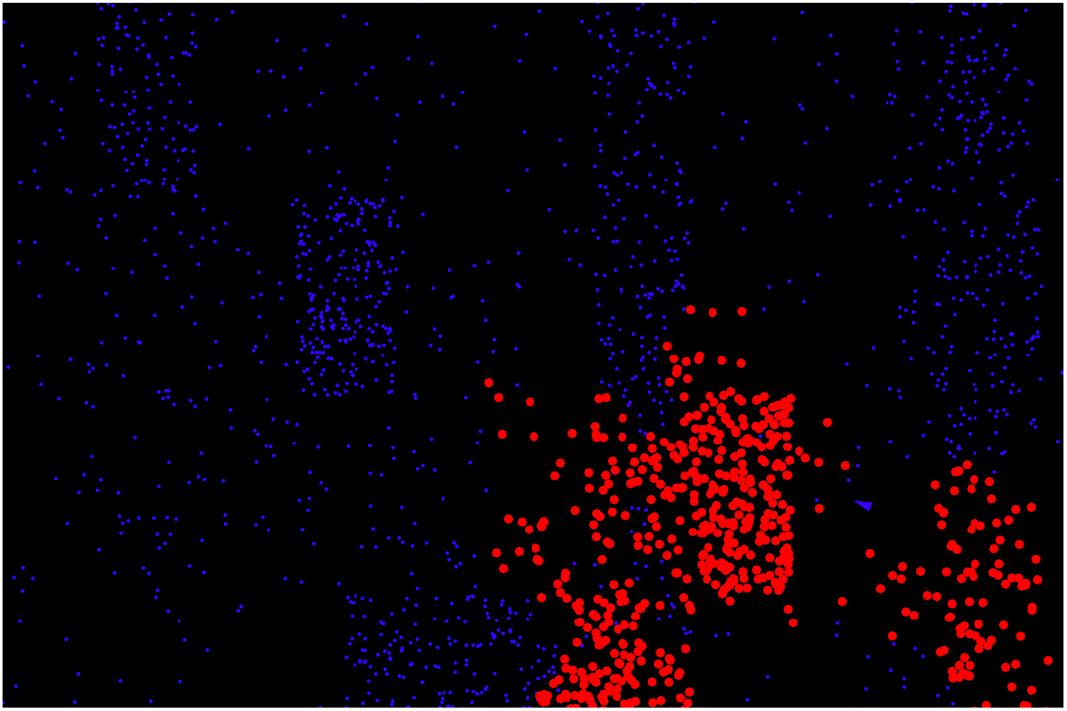
Simulation

Previously such models have been extensively studied using many parameters to govern the spread [4]. Also most human mobility models and realistic simulations which are needed for applications like SLAW [11] and SWIM [12] are based on Lévy Walks. There are also other compartmental models that typically forecast the rate of spread and behavioural count of the population (Such as the compartments of Susceptibles, Infectives and Recovered in the SIR model) [10, 7].

## 2 Simulation model

We take a two dimensional canvas of dimensions 1080×720 pixels characterised by blue dots indicative of population densities as shown in Fig. 2. This canvas may represent a continent, a country or a state where the scale invariant behaviour is observed, consecutively the blue dots have high density value if they represent a continent and low value if they represent a state. The population is highly concentrated in some regions and sparsely spread in other. A single autonomous carrier (blue triangle) moves around randomly controlled by ND or LD spanning across the canvas infecting and spreading the virus indicated by turning the blue to red dots. There maybe no direct physical significance to this carrier since the virus can spread by many different means. As the autonomous carrier moves along in a direction it has a perceptive radius of few pixels where it infects only the blue dots within a definite boundary surrounding it. The perceptive radius imitates the realistic situation where a certain carrier can only infect a specific region around. We run multiple iterations of a simulation with same parameters for different stable distributions to observe how the spread varies. We obtain the time taken by the autonomous carrier to infect fifty percent of the population and indicate it by *T*_50_, whereas the mean time for different iterations as 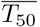. They are measured in seconds (s) where the wall time for each iteration is approximately equal to *T*_50_.

### 2.1 Comparing simulations of LD and ND

We initially observed which stable distribution is more ideal for the spread. The probability distribution function (PDF) for LD is described by

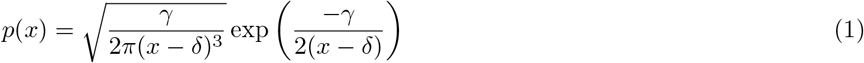

where *γ* is the scale parameter and *δ* is the location parameter. Similarly ND is described by

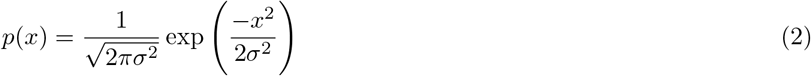

where *σ* is the standard deviation. *x* represents random values in the distribution with probability *p*(*x*) which are different lengths traversed by the carrier in pixels. For the LD shown in Fig. 3(a) where *γ* = 9.3 and *δ* = −3 we obtain 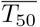 as 291 ± 21 s. We chose the parameters such that the peak for the PDF’s are comparable. Similarly for ND’s Normal1(N1) and Normal3(N3) whose standard deviation are *σ*(*N* 1) = 8 and *σ*(*N* 3) = 800 as shown in Figs. 3(a) and 3(b), we obtain 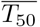 as 714 ± 173 s and 412 ± 57 s respectively. We take the standard deviation of N1 such that it is comparable with the perceptive radius of the autonomous carrier and the standard deviation of N3 is comparable with the size of the canvas. We observe that the spread in LW is more patchy but spans the entire region (Fig. 4(a)). Whereas in BW with low *σ* such as N1, the spread is more thorough and confined to particular regions (Fig. 4(b)). It takes longer *T*_50_ compared to LW and is more non deterministic. We also measure 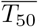 for other ND’s with standard deviation ranging between *σ*(*N* 1) and *σ*(*N* 3). *σ* of ND Normal2(N2) is the geometric mean of *σ*(N1) and *σ*(N3) as shown in Figs. 3(a), 3(b). For ND with *σ* ranging between *σ*(N1) and *σ*(N2), 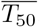 was obtained as 420 ± 64 s. 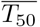 for N2 was obtained as 400 ± 53 s. Similarly for ND with *σ* ranging between *σ*(N2) and *σ*(N3), 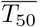 was obtained as 372 ± 37 s.

**Figure 3:**
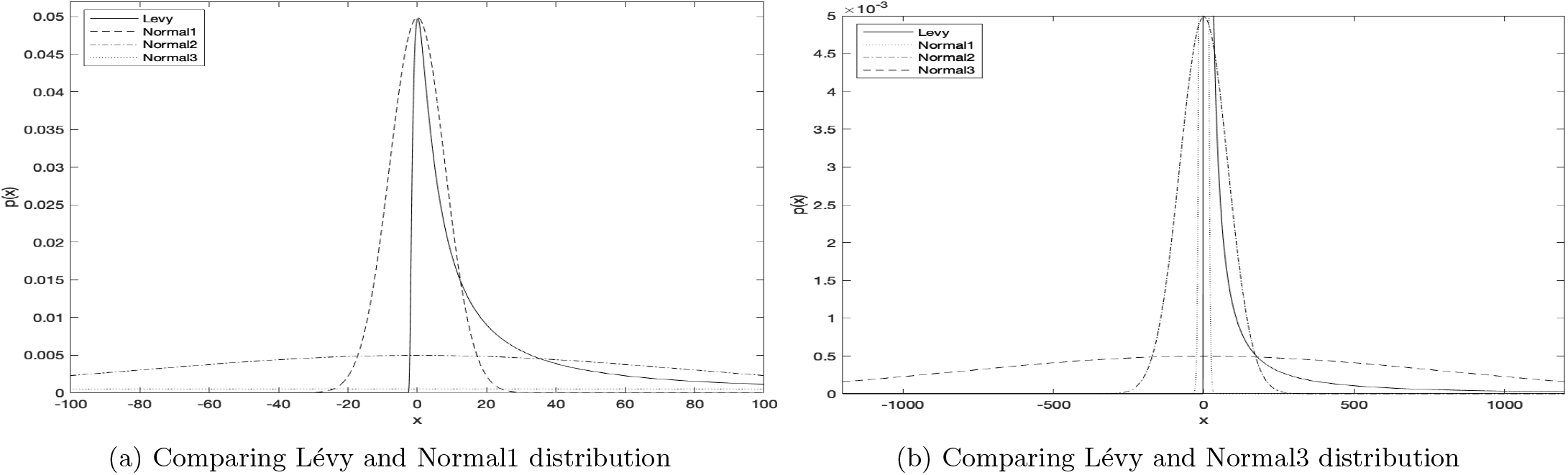
Comparing probability distributions

**Figure 4:**
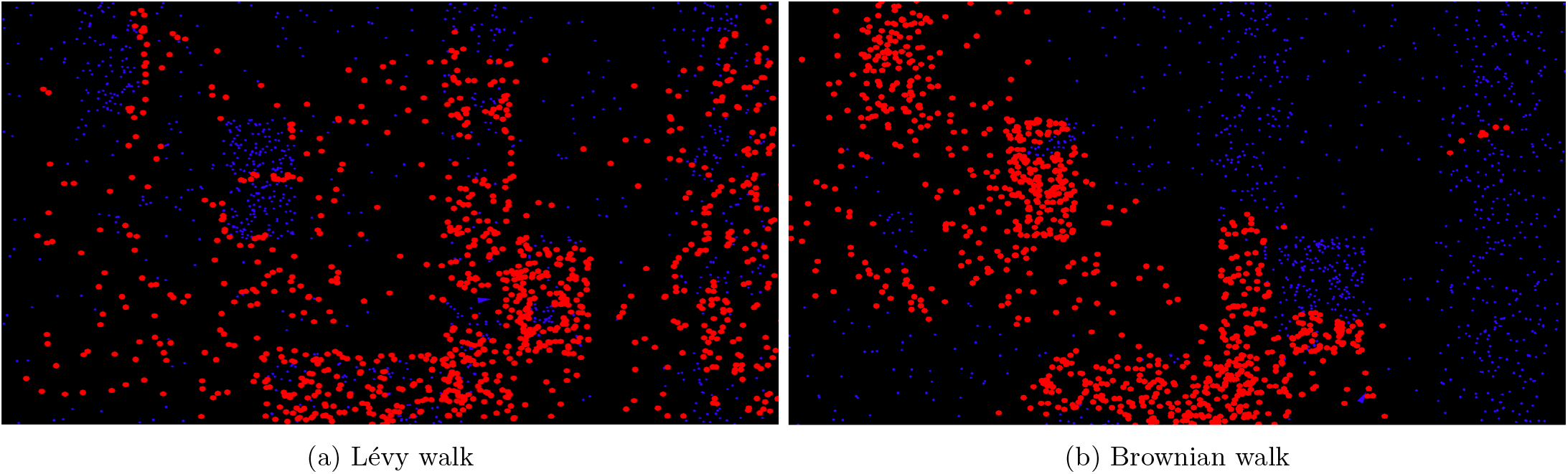
Simulations of random walk

### 2.2 Comparing simulations of LD with varying parameters *γ* and *δ*

We vary the parameters controlling LD, *γ* and *δ* to observe the change in spread by finding 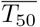 in each case. Location parameter *δ* determines where the peak of the probability distribution function lies and scale parameter *γ* determines the proportion of short steps and long steps. To study this wide range of parameter space faster simulation rates were used since we run around 50 hours of simulation. This may affect the accuracy of measuring 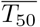 but a qualitative study can be made. We display 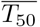 for varying parameters by factor of 10 in Table 1. We display the region of parameter space only where significant change was observed.

**Table 1:** 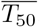 of LD with varying *γ* and *δ*

| $\delta \backslash \gamma$ | $10^{-6}$ | $10^{-3}$ | $10^{-2}$ | $10^{-1}$ | $10^0$ | $10^1$ |
| --- | --- | --- | --- | --- | --- | --- |
| $10^0$ | $60 \pm 10$ s | $60 \pm 7$ s | $57 \pm 8$ s | $58 \pm 7$ s | $58 \pm 9$ s | $54 \pm 7$ s |
| $10^1$ | $65 \pm 13$ s | $57 \pm 9$ s | $60 \pm 7$ s | $60 \pm 9$ s | $56 \pm 7$ s | $55 \pm 7$ s |
| $10^2$ | $50 \pm 7$ s | $51 \pm 5$ s | $49 \pm 6$ s | $49 \pm 6$ s | $51 \pm 6$ s | $51 \pm 7$ s |
| $10^3$ | $56 \pm 7$ s | $55 \pm 7$ s | $55 \pm 6$ s | $54 \pm 5$ s | $51 \pm 7$ s | $53 \pm 7$ s |

We do not find any deducible change in *T*_50_ by varying *δ* for each individual value of *γ*. For *δ* = 10^2^ the model dimensions are ideal where steps of lengths 10^2^ pixels are more prominent for uniform spread, without hitting the boundary and find 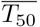 to be minimum. This is also observed when studying ND simulations with *σ* ranging between 80(N2) and 800(N3). Hitting the boundary or corners delays the spread since the carrier traverses through already infected regions and has to come back. Varying the *γ* for different *δ* we observe that smaller *γ* and *δ* values delay the spread where too many short steps take place compared to long steps. Long steps are necessary to cover the wide region and also *T*_50_ becomes more non deterministic in this parameter space. If the carrier is stuck taking many short steps in a region of highly crowded blue dots *T*_50_ value becomes low, whereas if stuck in a sparse region *T*_50_ is high. But also too many long steps in the region of higher *γ* and *δ* values makes the spread slower than ideal, with similar behaviour not affecting *T*_50_. We displayed newly infected blue dots per unit time of different iterations for *γ* = 10^−6^ in Fig. 5(a) and for *γ* = 10^1^ in Fig. 5(b). To reduce the boundary effects we chose *δ* = 10^2^ and observed how the spread varies for different *γ*. For lower *γ* value we observe a steady decline whereas for the higher *γ* value we observed fall and increase in the newly infected blue dots per unit time.

**Figure 5:**
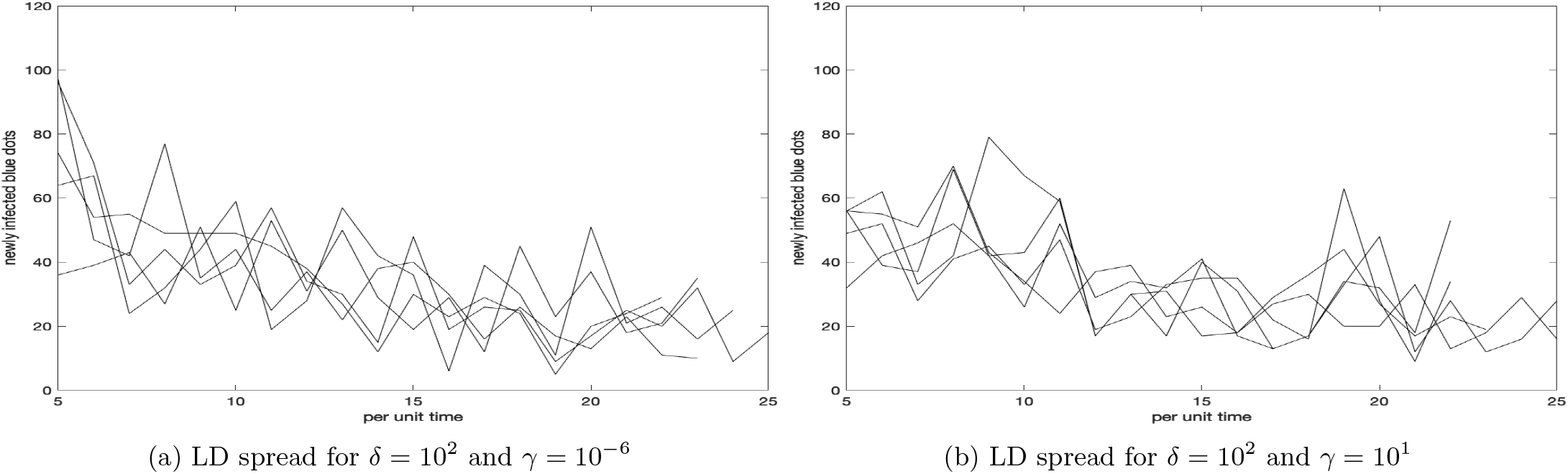
Newly infected blue dots per unit time for different iterations

### 2.3 Comparing simulations of log-normal distribution

Though a log-normal distribution has similar characteristics with LD and ND in the aspects that it can be expressed by sum of arbitary number of IID’s [19], it is not a stable distribution [5]. However since we are interested in such distributions whose normalized sum can reproduce the original distribution, we considered running simulations of same. Recent evidence suggests that individual modes of human mobility are based on log-normal distribution [20]. Log-normal distribution is described by PDF

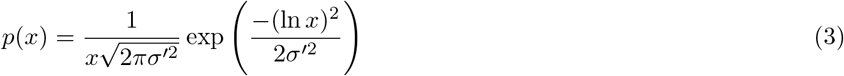

and we chose parameter *σ*′ such that the distribution is relatable to LD with heavy-tailed characteristics. For the same simulation rates used in Sec. 2.2 we do not observe much difference by varying *σ*′ ≥ 10 where we obtain 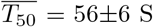, 56±8 Sand 57±6 S for *σ*′ = 10, 100 and 1000, respectively. Here the long steps are dominating which are essential for the spread. For *σ*′ < 10 we observe the PDF narrowed with limiting the number of long steps where we obtain 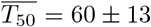 for *σ*′ = 1.

## 3 Conclusion

Initially we analysed that for our model, Lévy walk is the most ideal movement pattern for the spread. This may be related to the principle of least effort in the context of Zipf distribution [21], which have similar long-range correlations [18, 2]. It can perhaps be seen as by following Lévy walk the autonomous carrier takes the least effort to achieve maximum spread compared to Brownian walk. Further we studied how the parameters in Lévy and log-normal distributions affect the spread and how the behaviour of the spread varies. To consolidate, we implemented random variables from scale invariant continuous distributions as random events causing movement of a virus to study how the spread changes. Recently scale invariant discrete structures like fractals were implemented to study the spread of COVID-19[13]. Implementing such self-similarity into other mathematical models used to study the epidemic may lead to new insights. Future studies can be tried using more realistic parameters [4, 7] such as restricting the movement in infected zones, implementing multiple carriers and varying the dimensions of the simulations which could imitate real world scenario.

## References

[1] D. Brockmann, L. Hufnagel, and T. Geisel. The scaling laws of human travel. Nature, 439(7075):462–465, Jan 2006.

[2] Andras Czirók, Rosario N. Mantegna, Shlomo Havlin, and H. Eugene Stanley. Correlations in binary sequences and a generalized zipf analysis. Phys. Rev. E, 52:446–452, Jul 1995.

[3] Ensheng Dong, Hongru Du, and Lauren Gardner. An interactive web-based dashboard to track covid-19 in real time. The Lancet. Infectious diseases, 20(5):533–534, May 2020.

[4] Abdou Moutalab Fofana and Amy Hurford. Mechanistic movement models to understand epidemic spread. Philosophical Transactions of the Royal Society B: Biological Sciences, 372(1719):20160086, 2017.

[5] Xin Gao, Hong Xu, and Dong Ye. Asymptotic behavior of tail density for sum of correlated lognormal variables. International Journal of Mathematics and Mathematical Sciences, 2009:630857, Aug 2009.

[6] Marta C. González, César A. Hidalgo, and Albert-László Barabási. Understanding individual human mobility patterns. Nature, 453(7196):779–782, Jun 2008.

[7] Herbert W. Hethcote. The mathematics of infectious diseases. SIAM Review, 42(4):599–653, 2000.

[8] Mehran Kardar. Statistical Physics of Particles. Cambridge University Press, 2007.

[9] Christopher T Kello, Gordon D A Brown, Ramon Ferrer-I-Cancho, John G Holden, Klaus Linkenkaer-Hansen, Theo Rhodes, and Guy C Van Orden. Scaling laws in cognitive sciences. Trends in cognitive sciences, 14(5):223–232, May 2010.

[10] W. O. Kermack, and A. G. McKendrick,. A contribution to the mathematical theory of epidemics. Proceedings of the Royal Society of London. Series A, Containing Papers of a Mathematical and Physical Character, 115(772):700–721, 1927.

[11] K. Lee, S. Hong, S. J. Kim, I. Rhee, and S. Chong. Slaw: Self-similar least-action human walk. IEEE/ACM Transactions on Networking, 20(2):515–529, 2012.

[12] Alessandro Mei and Julinda Stefa. Swim: A simple model to generate small mobile worlds, 2009.

[13] Cristina-Maria Păacurar and Bogdan-Radu Necula. An analysis of covid-19 spread based on fractal interpolation and fractal dimension. Chaos, Solitons and Fractals, 139:110073, 2020.

[14] Andy M. Reynolds. Current status and future directions of lévy walk research. Biology Open, 7(1), 2018.

[15] C. W. Reynolds. Steering behaviors for autonomous characters. in Proceedings of the 1999 Game Developer’s Conference, 1999.

[16] I. Rhee, M. Shin, S. Hong, K. Lee, S. J. Kim, and S. Chong. On the levy-walk nature of human mobility. IEEE/ACM Transactions on Networking, 19(3):630–643, 2011.

[17] D. Shiffman. The nature of code.., 2012.

[18] Michael F. Shlesinger, George M. Zaslavsky, and Joseph Klafter. Strange kinetics. Nature, 363(6424):31–37, May 1993.

[19] Olof Thorin. On the infinite divisibility of the lognormal distribution. Scandinavian Actuarial Journal, 1977(3):121–148, 1977.

[20] Kai Zhao, Mirco Musolesi, Pan Hui, Weixiong Rao, and Sasu Tarkoma. Explaining the power-law distribution of human mobility through transportationmodality decomposition. Scientific Reports, 5(1):9136, Mar 2015.

[21] George Kingsley Zipf. Human Behavior and the Principle of Least Effort: An Introduction to Human Ecology. Addison-Wesley Press Inc., New York, 1949.

